# A comprehensive *in vivo* screen of yeast farnesyltransferase activity reveals broad reactivity across a majority of CXXX sequences

**DOI:** 10.1101/2023.02.06.527295

**Authors:** June H. Kim, Emily R. Hildebrandt, Anushka Sarkar, Wayland Yeung, La Ryel A. Waldon, Natarajan Kannan, Walter K. Schmidt

## Abstract

The current understanding of farnesyltransferase (FTase) specificity was pioneered through investigations of reporters like Ras and Ras-related proteins that possess a C-terminal CaaX motif that consists of 4 amino acid residues: Cysteine – aliphatic_1_ – aliphatic_2_ – variable (X). These studies led to the finding that proteins with the CaaX motif are subject to a 3-step post-translational modification pathway involving farnesylation, proteolysis, and carboxylmethylation. Emerging evidence indicates, however, that FTase can farnesylate sequences outside the CaaX motif and that these sequences do not undergo the canonical 3-step pathway. In this work, we report a comprehensive evaluation of all possible CXXX sequences as FTase targets using the reporter Ydj1, an Hsp40 chaperone that only requires farnesylation for its activity. Our genetic and high throughput sequencing approach reveals an unprecedented profile of sequences that yeast FTase can recognize *in vivo*, which effectively expands the potential target space of FTase within the yeast proteome. We also document that yeast FTase specificity is majorly influenced by restrictive amino acids at a_2_ and X positions as opposed to the resemblance of CaaX motif as previously regarded. This first complete evaluation of CXXX space expands the complexity of protein isoprenylation and marks a key step forward in understanding the potential scope of targets for this isoprenylation pathway.

## Introduction

Isoprenylation is a post-translational modification (PTM) catalyzed by several isoprenylation enzymes (i.e., FTase, GGTase-I, GGTase-II, GGTase-III). This PTM enhances protein association with cellular membranes or strengthens protein-protein interactions (Lane and Beese 2006, Leung, Baron et al. 2007, Tate, Kalesh et al. 2015, Wang and Casey 2016). Collectively, isoprenylated proteins assist in cellular signaling, cell cycle regulation, development, aging, and various other biological processes (Greenwood, Steinman et al. 2006, Palsuledesai and Distefano 2015, Jeong, Suazo et al. 2018).

Ras signaling GTPases are often cited as classical examples of isoprenylated proteins (Willumsen, Christensen et al. 1984, Wright and Philips 2006, Zhou, Wiener et al. 2016). Ras isoprenylation is catalyzed by farnesyltransferase (FTase), which acts upon a C-terminal CaaX motif consisting of 4 amino acid residues: Cysteine – aliphatic_1_ – aliphatic_2_ – variable (X). FTase appends the C15 isoprenoid lipid donated by farnesyl pyrophosphate to the CaaX cysteine via a thioether linkage. After farnesylation, Ras undergoes the coupled modifications of endoproteolysis to remove the -aaX portion of the motif, followed by carboxylmethylation of the farnesylated cysteine that becomes the new C-terminal residue. These PTMs regulate Ras plasma membrane localization and function, and have been extensively studied due to the significance of Ras GTPases in human disease such as cancer (Tamanoi 2011, Cox, Der et al. 2015, Hobbs, Der et al. 2016). As a result, previous Ras-based investigations have led to the general model that the three CaaX PTMs (i.e., isoprenylation, proteolysis, methylation) commonly occur across a range of proteins that are collectively referred to as CaaX proteins. CaaX PTMs exist in all eukaryotic species, and the impact of these PTMs on CaaX protein biology are highly conserved across model systems (Omer, Kral et al. 1993, Cazzanelli, Pereira et al. 2018, Ravishankar, Hildebrandt et al. 2023). This is especially evident in the conserved similarities beween and mammalian and yeast FTase structure and function (Kohl, Diehl et al. 1991, Gomez, Goodman et al. 1993, Omer, Kral et al. 1993).

Not all farnesylated proteins undergo all three CaaX PTMs. An example is the *Saccharomyces cerevisiae* Hsp40 chaperone Ydj1 (*Sc*Ydj1) that is farnesylated on its C-terminal CASQ sequence but not subjected to endoproteolysis or carboxylmethylation. Farnesylation of Ydj1 is required for optimal yeast growth at elevated temperatures (>37 °C) and subjecting Ydj1 to all three CaaX PTMs negatively impacts this Ydj1-dependent thermotolerance growth phenotype (Caplan et al., 1992, Hildebrandt et al., 2016). We have defined this non-canonical pathway leading to a farnesylation-only PTM as the “shunt” farnesylation pathway (**Figure 1**). The observation of the shunt pathway raises the question of whether the previous use of canonically-modified CaaX protein reporters (e.g., Ras, **a**-factor), which are subject to the additional constraints of proteolysis and carboxylmethylation, has limited the breadth of sequences that can be identified as FTase substrates.

**Figure 1.**
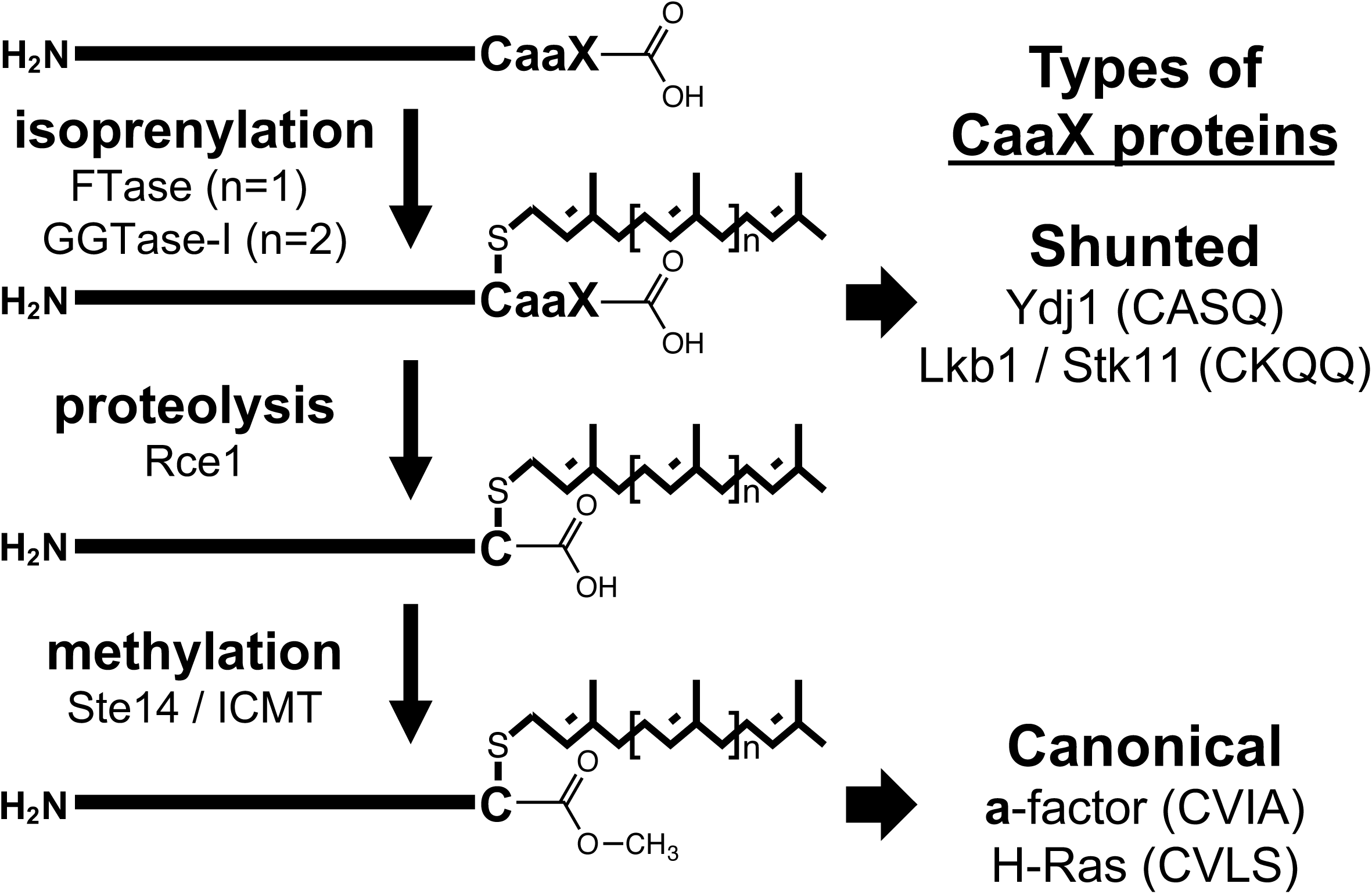
The post-translational isoprenylation pathway. Ydj1 and Lkb1/STK11 have non-canonical CaaX motifs that follow an isoprenylation-only pathway (i.e., shunted). Ras and **a**-factor have canonical CaaX motifs and are subject to the superposition of 3 modifications.

Many past investigations have attempted to probe FTase specificity. Initially, the screening of FTase substrates was explored *in vitro* on a case-by-case basis using recombinant CaaX proteins and synthetic peptides with [^3^H]-FPP or [^3^H]-GGPP, which was inconvenient and labor intensive for large scale studies (Moores, Schaber et al. 1991, Caplin, Hettich et al. 1994). *In vitro* investigations using peptide libraries and metabolic labeling have extended this effort but have been limited in scope and cost-prohibitive, making it difficult to conduct systematic studies of the 8000 CXXX sequence space (Reiss, Stradley et al. 1991, Boutin, Marande et al. 1999, Krzysiak, Scott et al. 2007, Krzysiak, Aditya et al. 2010, Wang, Dozier et al. 2014, Tate, Kalesh et al. 2015, Suazo, Schaber et al. 2016, Storck, Morales-Sanfrutos et al. 2019). While *in silico* prediction models based on structural analysis of mammalian FTase are available and have potential to define isoprenylatable sequences, these models often fail at predicting farnesylated proteins with non-canonical CaaX motifs such as *Sc*Ydj1 (CASQ), *Hs*DNAJA2 (CAHQ), *Sc*Pex19 (CKQQ), *Hs*Lkb1 (CKQQ) and *Sc*Nap1 (CKQS) and often lack orthologous *in vivo* reporter data to validate predictions (Collins, Reoma et al. 2000, Sapkota, Kieloch et al. 2001, Reid, Terry et al. 2004, Maurer-Stroh and Eisenhaber 2005, Lane and Beese 2006, Maurer-Stroh, Koranda et al. 2007, London, Lamphear et al. 2011, Berger, Yeung et al. 2022). Most of the *in vivo* reporters used to probe CaaX space have relied on CaaX protein reporters requiring 3-step modification and the superposition of 3 enzyme specificities for their activities (Boyartchuk, Ashby et al. 1997, Stein, Kubala et al. 2015). Moreover, recent reports demonstrating FTase activity on shortened and extended sequences highlight the flexibility of CaaX substrate lengths, additionally complicating the recognized model of FTase CaaX specificity (Ashok, Hildebrandt et al. 2020, Schey, Buttery et al. 2021).

The reliance of previous studies on canonical protein reporters and incomplete peptide reporter sets, in conjunction with our shunt pathway observations, led us to hypothesize that the full scope of FTase targets remains unknown. To address this gap in knowledge, we developed *Sc*Ydj1 as a genetic reporter to elucidate the specificity of the yeast FTase across all 8000 CXXX sequences. Our results indicate that yeast FTase has a much broader target specificity than previously defined by the canonical CaaX motif.

## Materials and Methods

### Yeast strains

Strains used in this study are listed in **Table 1**. All plasmid-transformations into yeast were performed using a lithium acetate-based transformation procedure unless otherwise stated (Elble 1992).

**Table 1.** Yeast strains used in this study.

| Identifier | Genotype | Reference |
| --- | --- | --- |
| BY4741 | *MATa his3Δ1 leu2Δ0 met15Δ0 ura3Δ0* | (Brachmann, Davies et al. 1998) |
| yWS304 | *MATa his3Δ1 leu2Δ0 met15Δ0 ura3Δ0 ydj1Δ::KAN<sup>R</sup>* | (Giaever, Chu et al. 2002) |
| yWS1632 | *MATa his3Δ1 leu2Δ0 met15Δ0 ura3Δ0 ram1::KAN<sup>R</sup>* | (Giaever, Chu et al. 2002) |
| yWS2542 | *MATa his3Δ1 leu2Δ0 met15Δ0 ura3Δ0 ydj1Δ::NAT<sup>R</sup> ram1Δ::KAN<sup>R</sup>* | (Berger, Kim et al. 2018) |
| yWS2544 | *MATa his3Δ1 leu2Δ0 met15Δ0 ura3Δ0 ydj1Δ::NAT<sup>R</sup>* | (Berger, Kim et al. 2018) |

### Plasmid construction for Ydj1-CKQx variants and His-Nap1

Plasmids used in this study are listed in **Table 2**. Newly created plasmids needed for this study were constructed using PCR-directed recombinational cloning consistent with previously reported methods (Oldenburg, Vo et al. 1997, Hildebrandt, Cheng et al. 2016, Berger, Kim et al. 2018). For Ydj1-CKQX plasmids, pWS1132 that was linearized with *Nhe1* was co-transformed into yWS304 along with a PCR-derived DNA fragment encoding a new CaaX sequence. For the His-Nap1 plasmid (pWS1474), pRS316 that was linearized with *BamHI*, *XbaI* and *XhoI* was con-transformed into BY4741 along with a PCR derived DNA fragment encoding the *NAP1* gene. The resultant Nap1 plasmid (pWS1318) was additionally modified to encode an octa-His tag at the 5° end of the *NAP1* gene; the tag was introduced by recombination using a PCR derived fragment and pWS1318 that had been linearized with *BsaBI*. In all cases, candidate plasmids recovered from yeast surviving appropriate genetic selection were evaluated by both restriction digest and DNA sequence analyses (GENEWIZ / Azenta Life Sciences, South Plainfield, NJ; Eurofins Genomics, Louisville, KY).

**Table 2.** Plasmids used in this study.

| Identifier | Genotype | Reference |
| --- | --- | --- |
| pRS316 | *CEN URA3* | (Sikorski and Hieter 1989) |
| pWS942 | *CEN URA3 YDJ1* | (Hildebrandt, Cheng et al. 2016) |
| pWS1132 | *CEN URA3 YDJ1-SASQ* | (Hildebrandt, Cheng et al. 2016) |
| pWS1286 | *CEN URA3 YDJ1-CVIA* | (Hildebrandt, Cheng et al. 2016) |
| pWS1318 | *CEN URA3 NAP1* | This study |
| pWS1407 | *CEN URA3 YDJ1-CKQQ* | This study |
| pWS1411 | *CEN URA3 YDJ1-CKQS* | (Berger, Yeung et al. 2022) |
| pWS1474 | *CEN URA3 His8-NAP1* | This study |
| pWS1563 | *CEN URA3 YDJ1-SKQQ* | This study |
| pWS1728 | *CEN URA3 YDJ1-BsrGI-SASQ* | This study |
| pWS2083 | *CEN URA3 YDJ1-CKQA* | This study |
| pWS2017 | *CEN URA3 YDJ1-CKQC* | This study |
| pWS2018 | *CEN URA3 YDJ1-CKQD* | This study |
| pWS2019 | *CEN URA3 YDJ1-CKQE* | This study |
| pWS2020 | *CEN URA3 YDJ1-CKQF* | This study |
| pWS2021 | *CEN URA3 YDJ1-CKQG* | (Berger, Yeung et al. 2022) |
| pWS2022 | *CEN URA3 YDJ1-CKQH* | (Berger, Yeung et al. 2022) |
| pWS2023 | *CEN URA3 YDJ1-CKQI* | This study |
| pWS2024 | *CEN URA3 YDJ1-CKQK* | This study |
| pWS2025 | *CEN URA3 YDJ1-CKQL* | (Berger, Yeung et al. 2022) |
| pWS2026 | *CEN URA3 YDJ1-CKQM* | This study |
| pWS2027 | *CEN URA3 YDJ1-CKQN* | This study |
| pWS2028 | *CEN URA3 YDJ1-CKQP* | This study |
| pWS2029 | *CEN URA3 YDJ1-CKQR* | This study |
| pWS2030 | *CEN URA3 YDJ1-CKQT* | This study |
| pWS2031 | *CEN URA3 YDJ1-CKQV* | This study |
| pWS2033 | *CEN URA3 YDJ1-CKQW* | This study |

### Plasmid construction for NGS library

The *YDJ1-CXXX* plasmid library (pWS1775) was designed to represent all 8000 “XXX” amino acid combinations where each amino acid is represented by a single unique codon by using trimer phosphoroamidites, aka “Trimer 20”, incorporated into the mutagenic oligos (Integrated DNA Technologies). To create the library, pWS1132 was first modified to introduce a silent unique *BsrGI* site directly 5° to the SASQ encoding sequence using oligo oWS1330 (5°-AACTATGATTCCGATGAAGAAGAACAAGGTGGCGAAGGTGTACAATCTGCATCTCAATGATTTTC), creating pWS1728 (*CEN URA3 YDJ1*-*BsrGI-SASQ*). Next, PCR-derived DNA fragments were generated by pairing oWS1359 (5°-TATGATTCCGATGAAGAAGAACAAGGTGGCGAAGGTGTACAATGT-iTriMix20-iTriMix20-iTriMix20-TGATTTTCTTGATAAAAAAAGATC-3°) and oWS1308 (5°-CAGCATATAATCCCTGCTTTA-3°) and pWS1728 template using a high-fidelity Q5 polymerase (New England Biolabs, Ipswich, MA). The PCR products were purified (Omega E.Z.N.A Cycle Pure kit, Omega Bio-tek Inc., Norcross GA), digested with BsrGI and AflII, re-purified, and ligated into pWS1728 (BsrGI and AflII digested) using high concentration T4 DNA ligase (NEB M0202T). Ligations were transformed into DH5⍺ *E. coli* prepared using Z-Competent cell kit (G-Biosciences, St. Louis, MO) and plated to achieve approximately 10^4^ colonies per plate. In total, 606,800 colonies, each representing a unique clone, were pooled into a single cell pellet, followed by DNA purification (Omega E.Z.N.A Midiprep kit, Omega Bio-tek Inc., Norcross GA).

### Preparation of naïve yeast library containing Ydj1-CXXX variants

The *E. coli*-derived Ydj1-CXXX plasmid library was transformed into yWS304 (*ydj1*Δ) via Frozen-EZ Yeast transformation II Kit by Zymo Research (Irvine, CA) according to manufacturer’s instructions such that over 3 million individual colonies were recovered across multiple SC-Uracil plates on the same day. The colonies were collected by gently washing the plates with liquid medium, and cells concentrated by centrifugation to remove excess supernatant. The collected cells were stored at −80 °C as 100 µL aliquot stocks in 15% glycerol at a concentration of ∼2×10^9^ cells per 1 mL. For all subsequent studies, freshly thawed stock vials were used and not re-frozen for repeated use.

### Thermotolerance selection of Ydj1-CXXX variants *en masse* and plasmid library recovery

An aliquot of the naïve yeast library expressing Ydj1-CXXX variants was thawed and used for each iteration of this assay. In brief, ∼582,000 cells were diluted into 40 mL of room temperature SC-Uracil liquid media, and this premix was divided across 4 test tubes where sets of 2 tubes were incubated at either permissive (25 °C) or restrictive temperature (37 °C). Each test tube containing diluted culture was rapidly thermally equilibrated using an appropriate temperature water bath for 30 minutes before being placed onto a rotating wheel in an incubator at the same temperature. The cultures were incubated for approximately 24 to 48 hours until A_600_ 2.0 was reached, cells were collected by centrifugation, and plasmids were isolated from the recovered cell populations using a commercial kit (OMEGA Bio-Tek E.Z.N.A. Yeast Miniprep kit) as per manufacturer’s instructions. The experiment was performed twice across two different days, with each condition performed in duplicate, for a total of 8 replicates per temperature condition.

### Next-generation sequencing and data analysis

The yeast plasmids recovered after thermoselection were subject to a shortened PCR with 15 cycles with oligonucleotide pairs oWS1408 (5°-GTCTCGTGGGCTCGGAGATGTGTATAAGAGACAGCAGCATATAATCCCTGCTTTA-3°) and oWS1409 (5°-TCGTCGGCAGCGTCAGATGTGTATAAGAGACAGTCCAGGGGTGGTGCAAACTATG-3°) using a high-fidelity Q5 polymerase (New England Biolabs, Ipswich, MA) to attach overhang sequences. The PCR products were cleaned (OMEGA Bio-Tek E.Z.N.A. Cycle Pure Kit), quantified (Synergy H1 Hybrid Multi-Mode Microplate Reader), and submitted to the Georgia Genomics Bioinformatics Center (GGBC, Athens, GA) for Illumina MiSeq library assembly and sequencing (single-paired 100 bp reads starting 38 bases upstream of the CXXX region of interest).

The NGS analysis yielded over 10 million total reads, with ∼96% of individual reads at or above the quality score of Q30. All 8 replicates from 25 °C yielded high-quality reads while only 7 replicates from 37 °C passed the quality cutoff. These 15 experimental samples, which represented about 3 million total reads, were carried forward for downstream analysis. In parallel, NGS was used to sequence 10 replicates of the *E. coli* plasmid library and 10 replicates of the plasmids extracted from the naïve yeast library prior to selection.

To assess the likelihood of farnesylation of Ydj1-CXXX variants, the reads within each replicate were first assessed to determine the number of occurrences for each unique CXXX sequence present. The occurrence of each CXXX sequence was summed across all replicates of the same temperature condition, then normalized to a frequency value. The latter was calculated by dividing the occurrence value for each unique CXXX by the total number of occurrences of all CXXX sequences (i.e., frequency = count_unique_/count_total_) for each of the 25 °C and 37 °C data sets. Lastly, the frequency value of each unique CXXX sequence was used to determine a unique enrichment factor (EF) score. This was calculated by dividing the frequency value of a unique CXXX sequence at restrictive temperature by that of the 25 °C condition (i.e., EF score = frequency_37 °C_/frequency_25 °C_). Some CXXX sequences were not recovered at 25 °C, yet present at higher temperatures (n=9; CFFM, CIFF, CILF, CIYM, CNWC, CTFA, CVFF, CVLW, CWIA). To account for such cases, a correction factor of +1 was applied across all frequency values at 25 °C so that 1 would be the denominator for CXXX sequences with 0 occurrences at 25 °C. We also applied the same correction factor of +1 to all frequency values at 37 °C for consistency.

### Weblogo sequence alignments

The top 5% (n=400) sequences for both Ydj1-based and Ras-based screens were evaluated by Weblogo (http://weblogo.berkeley.edu/logo.cgi) using a custom color scheme (Crooks, Hon et al. 2004). Cys was set to blue; polar charged amino acids were set to green (Asp, Arg, Glu, His, and Lys); polar uncharged residues were set to black (Asn, Gln, Ser, Thr, and Tyr); branched chain amino acids were set to red (Ile, Leu and Val); all other residues were set to purple (Ala, Gly, Met, Phe, Pro, and Trp).

### Heatmap analysis

CXXX variants were clustered into 20-member groups having shared amino acids pairs. An average EF value was determined for these groups to allow for contextual analyses exploring the relationships between a_1_ vs. a_2_, a_1_ vs. X, and a_2_ vs. X. The averages were then analyzed with Microsoft Excel version 16.65 using the Conditional Formatting, Color Scales (Green - Yellow - Red) function to produce three distinct heatmaps (HMs). The values reported within each cell of a heatmap represent the average EF value for each respective 20-member group. The EF scores associated with RRS data (Stein et al., 2015) were evaluated in the same manner.

To determine confidence intervals, the average HM values were averaged across each individual row and column of the heatmap, and the averages were used to determine a standard deviation for either the row or column set of values. A standard deviation calculation was next used to determine a 95% confidence interval that was in turn used to establish high and low cutoffs for identifying positive selection or negative restriction. A pattern was deemed significant when 18 or more of each 20-member set (i.e., >90%) were above or below the statistical cutoff.

### Likelihood of prenylation calculations for each CXXX variant

The averaged EF values from each of the 3 HMs were summed to produce a HM score. For example, to predict the likelihood of prenylation for CASQ, the averages of Cys-Ala-Ser-X (n=20 varied at the X position), Cys-Ala-X-Gln (n=20 varied at the a_2_ position), and Cys-X-Ser-Gln (n=20 varied at the a_1_ position) were first calculated, then these three HM values were summed to represent the HM score of CASQ.

### Ydj1-CXXX thermotolerance assay

The assay was performed as previously described (Hildebrandt, Cheng et al. 2016, Berger, Kim et al. 2018, Ashok, Hildebrandt et al. 2020). In brief, yeast cultures in SC-Uracil media were incubated at 25 °C until saturation, then serially diluted 10-fold in YPD before being pinned onto YPD solid medium. Plates were incubated for 2-4 days at 25 °C, 37 °C, and 39 °C prior to results being digitally scanned face down without lids using a Cannon flat-bed scanner (300 dpi; TIFF format). Scanned images were adjusted for consistency in image rotation, contrast, and size before being copied onto Microsoft PowerPoint version 16.65 for final figure assembly. The experiment was performed twice in duplicate.

### Gel-shift assay

The assay was performed as previously described (Hildebrandt, Cheng et al. 2016, Berger, Kim et al. 2018, Schey, Buttery et al. 2021). Whole cell lysates of mid log cells were prepared and separated by sodium dodecyl sulfate-polyacrylamide gel electrophoresis (SDS-PAGE; 6% stacking with 9.5% resolving gel) then transferred onto nitrocellulose. Blots were blocked with 5% milk then sequentially incubated with rabbit anti-Ydj1 primary antibody (courtesy of A. Caplan) and HRP-conjugated goat anti-rabbit secondary antibody (Kindle biosciences, Greenwich, CT). Fluorescence was detected using the KwikQuant Western Blot Detection Kit (Kindle Biosciences) and a KwikQuant Imager and as per manufacturer’s instructions. Protein bands were quantified using NIH ImageJ, and resulting values were used for calculating ratios for prenylated and non-prenylated bands. The blots containing yeast Nap1 were treated similarly but incubated with mouse anti-HIS primary antibody (Thermo Fisher Scientific, Waltham, MA) and HRP-conjugated sheep anti-mouse secondary antibody (Kindle biosciences, Greenwich, CT).

### Decision tree modeling

Sequence motif features were generated by one-hot encoding for each of the variable CaaX motif sites: a_1_, a_2_, and X. Additional binary features were included to describe whether the variable residue was within a given set of residues. To define these sets, we enumerated all possible amino acid combinations with a max set size of 5. In total, 65,097 features were considered. While building the decision tree classifier, entropy was used to evaluate the quality of potential splits. To determine more generalizable rules, trees were allowed a maximum depth of 3 and all nodes were required to have a minimum of 50 samples. This method was implemented using Scikit-learn (v0.22.2) using the DecisionTreeClassifier class.

## Results

### Many CXXX sequences sustain Ydj1-dependent growth of yeast at high temperature

Ydj1 follows an isoprenylation-only pathway in contrast to canonical reporters (i.e., Ras GTPases, **a**-factor) that undergo additional downstream modifications (Trueblood, Boyartchuk et al. 1997, Stein, Kubala et al. 2015) (**Figure 1**). As such, the use of Ydj1 as a reporter to probe yeast FTase specificity reduces the risk of introducing additional specificity bias from CaaX proteases. To evaluate the substrate scope of FTase, we adapted methods from a previous yeast study that utilized a Ras-based CXXX reporter (i.e., Ras Recruitment System (RRS)) along with competitive growth enrichment and next-generation sequencing (NGS) methods (Stein, Kubala et al. 2015). In our case, we investigated the ability of Ydj1-CXXX variants to sustain high temperature growth in a *ydj1Δ* genetic background (**Figure S1**). A plasmid library of Ydj1-CXXX variants was generated using single codons for each amino acid so that all 8000 CXXX sequences are represented in a relatively small library (see Material and Methods). Compared to traditional plasmid library construction relying on fully or partly degenerate oligonucleotides, a Trimer 20-based strategy was used with the aim of yielding a balanced library with respect to codon redundancy (i.e., no over-representation of amino acids with multiple codons) and no introduction of early stop codons, which effectively reduces the number of copies needed to reach statistical confidence for full coverage (Firth and Patrick 2008). While 110,500 independent clones were minimally required for statistical 100% coverage, our library contains 606,800 independent clones (i.e., 6x full coverage). By contrast, constructing a library with fully degenerate oligonucleotides to vary 3 amino acids would have required 3,200,000 independent clones for 100% coverage, leading to much higher labor and time costs for this study.

The Ydj1-CXXX plasmid library DNA was purified from *Escherichia coli* before being transformed into the *ydj1Δ* yeast strain to yield a naïve *ydj1Δ/YDJ1-CXXX* yeast library. Both the *E. coli* plasmid library and the naïve yeast library were confirmed by NGS to contain all potential Ydj1-CXXX variants (**File S1**). Ligation efficiency of vector alone was determined to be ∼0.3% relative to vector plus insert, indicating that a small portion of the library could encode *YDJ1-SASQ* (i.e., uncut parent plasmid). Consistently, *YDJ1-SASQ* was observed at frequencies of 0.065% and 0.071% in the *E. coli* and naïve yeast libraries, respectively. The frequencies of all CXXX sequences from the *E. coli* and naïve yeast libraries were also graphed, and no significant changes in the frequency profile between the two libraries were observed (**Figure S2**). This analysis further revealed that the libraries were not perfectly balanced such that the extremes exhibited a ∼10x range in abundance.

The naïve yeast library was propagated for approximately 8 generations at permissive (25 °C) and selective (37 °C) temperatures in liquid media until the culture was well saturated (A_600_ ∼2.0). The selective temperature condition was expected to enrich for farnesylated Ydj1-CXXX variants. NGS methods were then used to identify all the CXXX sequences present in each culture, from which frequencies were determined for each CXXX sequence in each temperature group. Similarities in experimental design allowed for direct comparison of Ydj1 and Ras-based data sets. In the RRS study, the likelihood of farnesylation was reported by an Enrichment Factor (RRS EF) that was defined by the frequency of a specific CXXX sequence occurring at 37 °C divided by its frequency at 25 °C (i.e., frequency_selective_/frequency_permissive_). A high RRS EF was interpreted as a high possibility of farnesylation. We performed a similar calculation for the Ydj1 data.

From our analysis, we observed that EFs for the Ydj1-based screen exhibited a narrower range (EF: 0.036 for CWWC to 13.898 for CVFF) relative to the RRS EFs for the Ras-based screen (RRS EF: 0.008 for CLRS to 70.667 for CYCM). We interpret these ranges to indicate that the majority of all CXXX sequences can support growth in the Ydj1-based screen, whereas a smaller number of sequences undergo higher enrichment during the thermoselection process in the Ras-based screen. The comparison also revealed that the EF profiles offered two distinct sequence landscapes, especially for farnesylated sequences that are well characterized: non-canonical CASQ (*Sc*Ydj1) and CKQQ (*Hs*STK11/Lkb1); canonical CVIA (*Sc* **a**-factor) and CVLS (*Hs*H-Ras). In the Ydj1 NGS-based screen, non-canonical sequences CASQ (EF: 1.608) and CKQQ (EF: 1.627) outperformed canonical CaaX sequences such as CVIA (EF: 0.406) and CVLS (EF: 0.494) (**Figure 2A**). By contrast, in the Ras-based screen, CASQ (RRS EF: 0.399) and CKQQ (RRS EF: 0.399) were significantly less enriched while CVIA (RRS EF: 10.124) and CVLS (RRS EF: 11.804) ranked amongst the top hits (**Figure 2B**). The EFs of all CXXX sequences resulting from the Ydj1 screen are reported in **File S2**.

**Figure 2.**
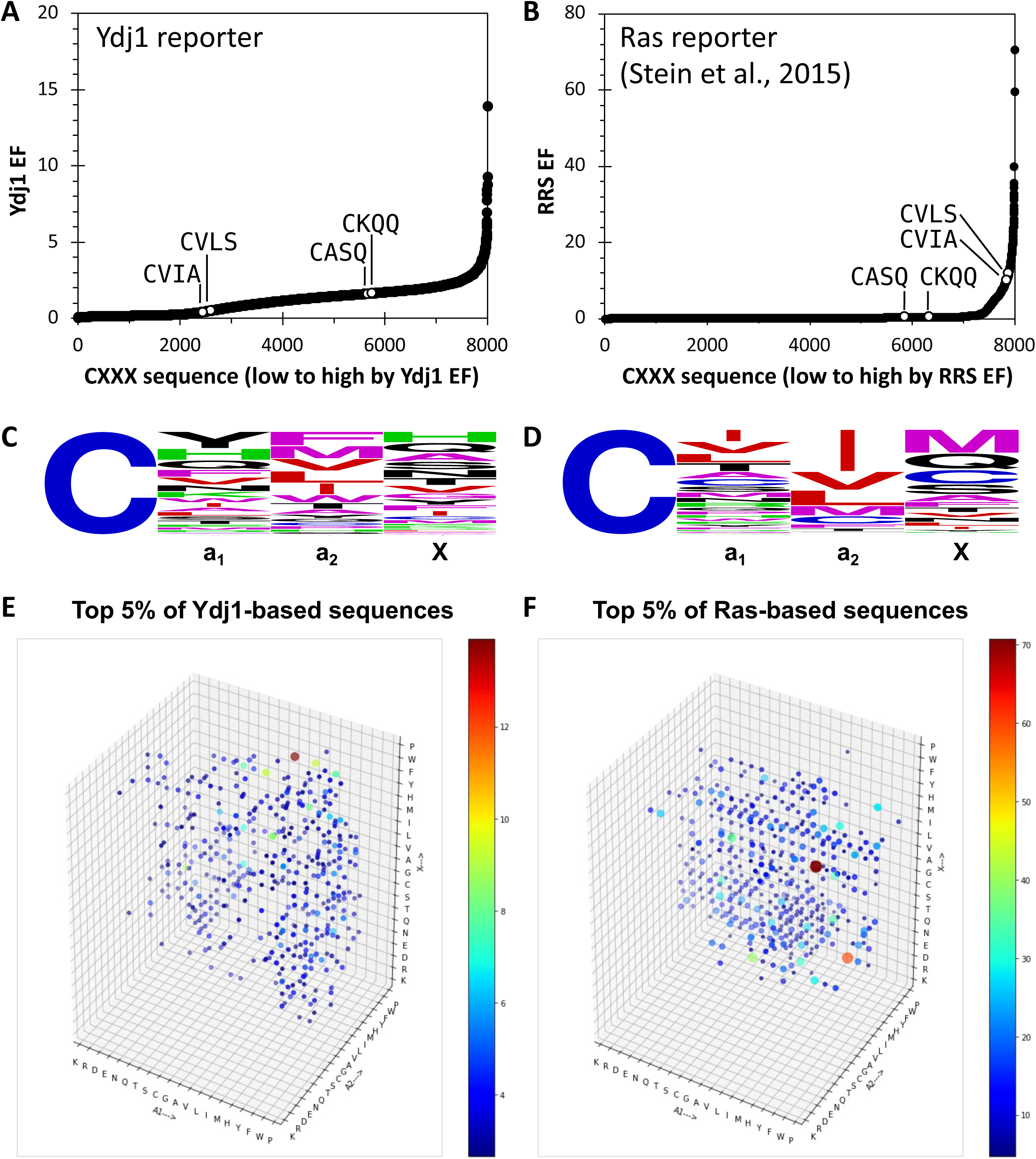
Comparison of results from the Ydj1-CXXX NGS screen and previously published Ras-CXXX NGS screen. Enrichment profiles of (**A**) Ydj1-based and (**B**) Ras-based screens offer two distinct sequence landscapes. Annotated controls: non-canonical *Sc*Ydj1 (CASQ) and *Hs*Lkb1/STK11 (CKQQ); canonical *Sc* **a**-factor (CVIA) and *Hs*H-Ras (CVLS). Weblogo frequency representation of top 5% (n=400) of hits for (**C**) Ydj1-based and (**D**) Ras-based screens. A 4D representation of the space occupied by top 5% of hits recovered from (**E**) Ydj1-based and (**F**) Ras-based screens. The size of the dot is the 4th dimension and corresponds to the enrichment factor of the CXXX motif within the respective screen.

Our observations suggest that the Ydj1-based screen best enriches non-canonical sequences whereas the Ras-based screen best enriches canonical sequences. The top 5% (n=400) of hits from the Ydj1 NGS-based assay did not exhibit an obvious consensus sequence (**Figure 2C**); however, the same number of top hits from the Ras-based screen were enriched with aliphatic amino acids at the a_1_ and a_2_ positions (**Figure 2D**). In the context of the Ras reporter, an aliphatic amino acid is especially prominent at the a_2_ position, which has been historically regarded as a requirement for FTase specificity. The top 5% of hits from both the Ydj1-based and Ras-based screens were also displayed as a 4D plot (**Figures 2E, F**; the size of the spot is the 4^th^ dimension and represents relative abundance). This analysis revealed that the top hits were more widely dispersed across CXXX sequence space in the Ydj1-based data relative to the Ras-based data, and that the most abundant sequences differed between the data sets. Analysis of the data as 3D plots, viewed along each of the 4D plot axes such that Lysine (K) is the nearest amino acid, provided additional details about the sequence space covered by each screen (**Figure S3**). In these 3D plots, the spots representing relative abundance are stacked behind each other, and the total number of spots occupying a particular node is not easily discerned. Although this arrangement makes it difficult to make detailed conclusions about the depth of coverage at each node (i.e., the number of amino acids present), it allows for clear conclusions about restrictions (i.e., amino acids that are not tolerated independent of context). From the perspective of a_1_, both data sets lack several amino acids at a_2_ (i.e., D, E, G, K, and R) and a single amino acid at X (i.e., R) (**Figures S3A, B**). Specific to the Ras-based data, additional amino acids were absent at a_2_ (H and Y) and X (K, P, and W) or less prevalent at a_2_ (i.e., A, P, Q, S, T, and W). From the perspective of a_2_, no amino acids were absent at the a_1_ position in both data sets, one amino acid was absent at the X position in both data sets (i.e., R), and several amino acids were absent at the X position within the Ras-based data set only (i.e., K, P, R, and W) (**Figures S3C, D**). From the perspective of X, no amino acids were absent at the a_1_ position, and the amino acids that were absent at a_2_ from the perspective of a_1_ were again identified, with one additional amino acid being absent (i.e., S) (**Figures S3E, F**). A general interpretation of these observations is that many amino acids can be accommodated at each position of the CXXX sequence independent of whether Ydj1 or Ras is the reporter, except for charged residues that are strictly not tolerated at a_2_ in either case. There also appear to be additional exceptions in the context of the Ras reporter at both the a_2_ and X position that likely reflect its need for additional modification beyond initial isoprenylation.

To further analyze our Ydj1 NGS-based data, we crosschecked CXXX sequences that were previously identified as being farnesylated through a limited scope genetic Ydj1-based Temperature Screen (YTS) (Berger, Kim et al. 2018). Mostly non-canonical sequences were identified in the earlier study (n=153). Many of these sequences exhibited high EFs in the present study (**Figure 3A**). When superposed on the Ydj1 EF profile, most YTS identified sequences were positioned in the 4th quartile (n=77) with fewer in the 3^rd^ quartile (n=67) and fewest in the 2^nd^ quartile (n=9). None of the YTS hits were present in the 1^st^ quartile. By contrast, superposition of the top hits from the Ras-based screen (n=496; RRS EF > 3; the cutoff for positive hits of the Ras-based screen) on the Ydj1 EF profile revealed positioning of sequences in the 2^nd^, 3^rd^, and 4^th^ quartiles of the EF profile (n=228, n=114, and n=153, respectively), with the fewest in the 1^st^ quartile (n=1) (**Figure 3B**). The few canonical sequences identified by YTS (**Figure 3A**, dashed box; n=15) remained within the range displayed by RRS sequences. Together, these results are fully consistent with our Ydj1 NGS-based method being useful for enriching non-canonical sequences that are farnesylated in addition to canonical sequences observed using the Ras reporter.

**Figure 3.**
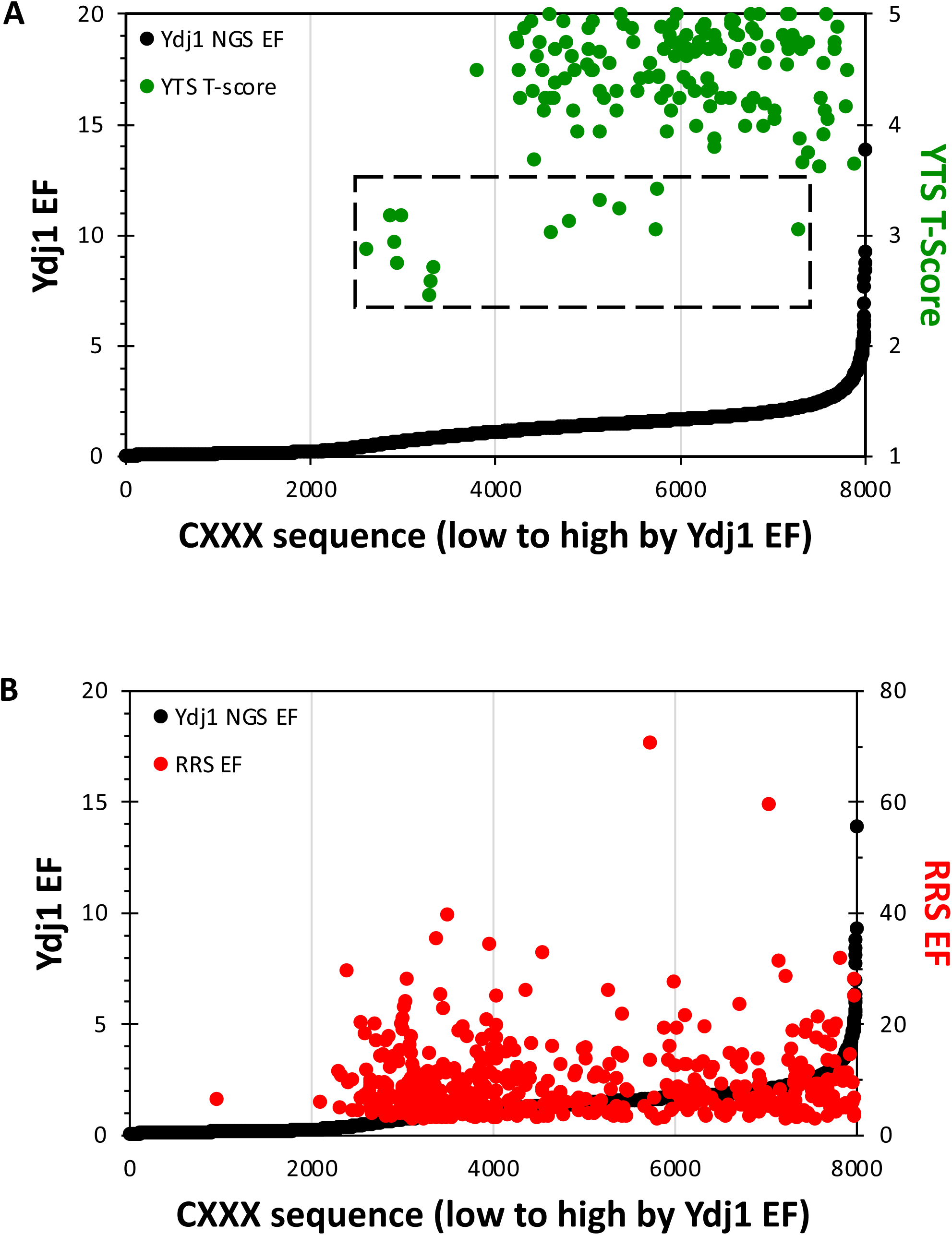
Relative performance of previously identified farnesylated CXXX sequences. Sequences previously identified in a (**A**) Ydj1-based genetic selection or a (**B**) Ras-based competitive growth assay were superimposed on the Ydj1 EF profile. Each sequence is represented as a point with an associated score where the second Y-axis refers to the scoring range used in these previous studies (YTS T-score - Ydj1-based Temperature Screen Thermotolerance Score; RRS EF – Ras Recruitment System Enrichment Factor). The limited number of canonical sequences that were identified by YTS are identified by a dashed box (n=15).

### Non-canonical CKQX sequences are targeted by FTase

Emerging evidence indicates that some CKQX sequences are farnesylated despite the presence of non-aliphatic amino acids at a_1_ and a_2_ positions. CKQQ farnesylation is well documented for human STK11/Lkb1, human Nap1L1 and yeast Pex19, whereas CKQS derived from yeast Nap1 is farnesylated in the context of the Ydj1 reporter (Collins, Reoma et al. 2000, Sapkota, Kieloch et al. 2001, Tate, Kalesh et al. 2015, Storck, Morales-Sanfrutos et al. 2019, Berger, Yeung et al. 2022). We additionally confirmed that farnesylation of CKQS occurs in the natural context of yeast Nap1 itself using a gel-shift assay that evaluates the mobility of farnesylated species by SDS-PAGE (**Figure S4A**). Of note, farnesylation has an opposite effect on Nap1 mobility when compared to Ydj1. Moreover, the ScanProsite tool (https://prosite.expasy.org/scanprosite/) identifies 5179 entries across UniProtKB/Swiss-Prot and UniProtKB/TrEMBL reference proteome sequences ending in CKQX across eukaryotes (**File S3**). The majority of these sequences (i.e., 65%) contain CKQQ or CKQS. Out of the 20 CKQX variants, most (n=17 positive hits) displayed a Ydj1 EF consistent with a high likelihood of farnesylation (**Figure 4A**). The remaining sequences (n=3) had low EFs, were well separated from the other CKQX sequences on the EF profile, and were expected to have a low likelihood of modification (i.e., negative hits). By comparison, none of the 20 CKQX sequences were predicted to be targets of FTase in the Ras-based EF profile (**Figure 4B**).

**Figure 4.**
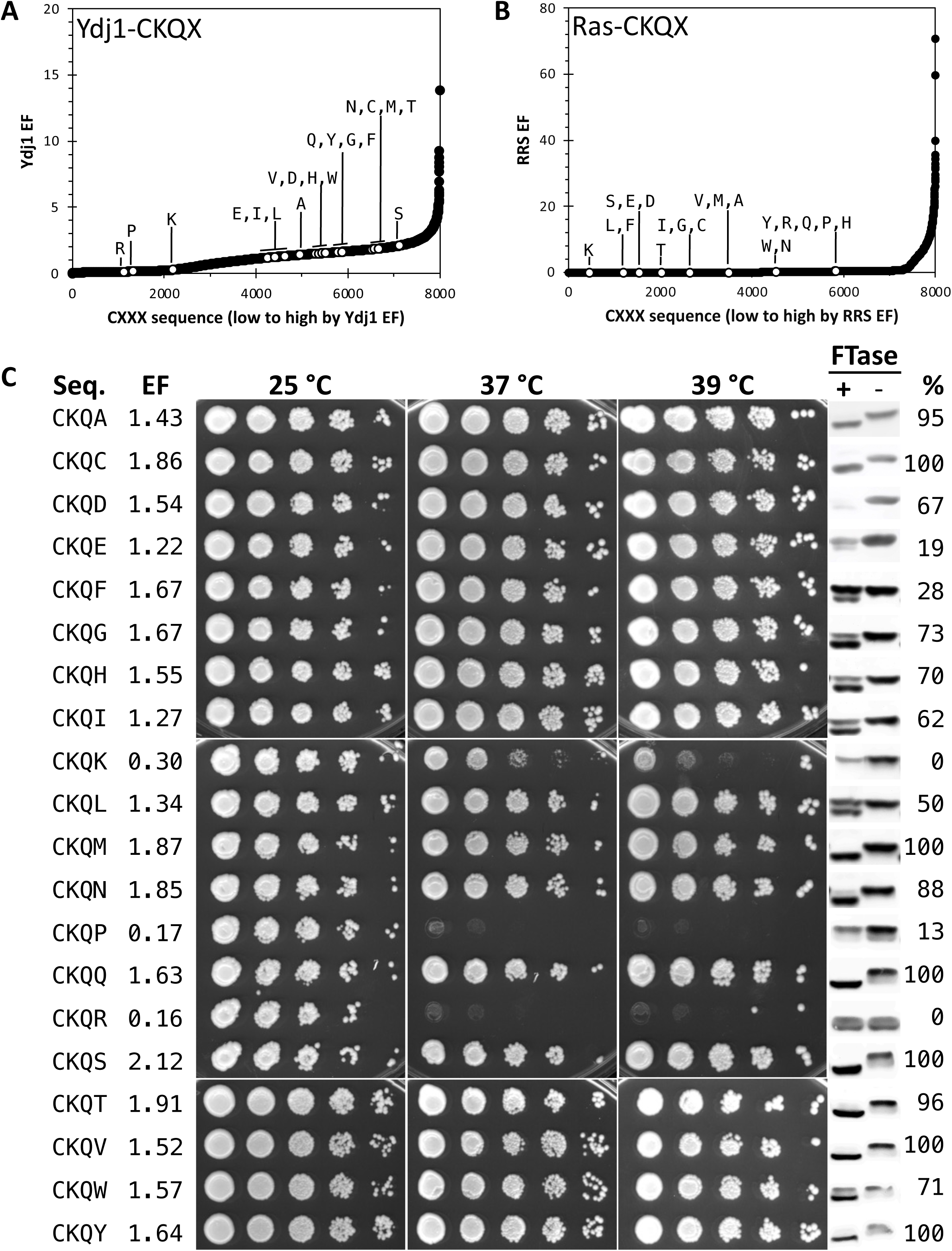
Evaluation of CKQX variants as FTase substrates. Enrichment profiles of CKQX sequences in (**A**) Ydj1-based and (**B**) Ras-based screens. **C**) Thermotolerance and gel-shift assays of Ydj1-CKQX variants provide supporting evidence towards the prenylation of non-canonical CKQX sequences by yeast FTase. Thermotolerance assays were performed by culturing yeast to the same density, applying a 10x serial dilution to the cultures, spotting dilution sets onto YPD rich media, and incubating at indicated temperatures. Notably, plate-based selection requires a slightly higher temperature to mimic the growth profile observed on liquid-based selection. The strains used were transformants of yWS304 containing the indicated Ydj1-CXXX variant. Gel shift assays were performed for the indicated Ydj1-CXXX variants in the presence and absence of FTase activity. Total yeast extracts were analyzed by SDS-PAGE and immunoblotting. The strains used were yWS2544 (+FTase) and yWS2542 (*ram1*Δ; - FTase. Abbreviations: Seq = Sequence, EF = Enrichment Factor, % = percent prenylation.

The farnesylation status of Ydj1-CaaX variants were monitored by thermotolerance and gel-shift assays (Caplan, Tsai et al. 1992, Hildebrandt, Cheng et al. 2016). The thermotolerance assay confirmed the 17 CKQX positive hits to have growth profiles similar to that of wildtype Ydj1, whereas the negative hits (n=3) had a growth profile consistent with unmodified Ydj1 (**Figure 4C, Figure S4B**). In strong agreement with the EFs and thermotolerance profiles, a prenyl-dependent gel-shift was observed for the 17 positive Ydj1-CKQX hits and no shift observed for the three negative hits. It should be noted that CKQR displayed a doublet gel pattern in both the presence and absence of FTase. The reason for this doublet pattern is unknown, however, the pattern was identical with and without FTase, so CKQR was deemed as having 0% prenylation.

### Statistical analyses reveal yeast FTase specificity

Our evaluation of genetic and biochemical data suggested that FTase is not limited to target sequences having aliphatic amino acids at a_1_ and a_2_. To identify factors that could better indicate target specificity, we assessed the likelihood of prenylation for sets of CaaX motifs that shared the same context, differing only at a single amino acid position. Using heatmap analysis, we sought to identify potential patterns marked by positive selection (colored in green) or negative restriction (colored in red) across a_1_, a_2_, and X positions in both Ydj1-based and Ras-based results (**Figure 5**). These patterns were identified as instances where the HM values of 18 or more of each 20-member set (i.e., >90% of EF scores) were well above or below the statistical average for all values (n=400) within a heatmap (**File S4**).

**Figure 5.**
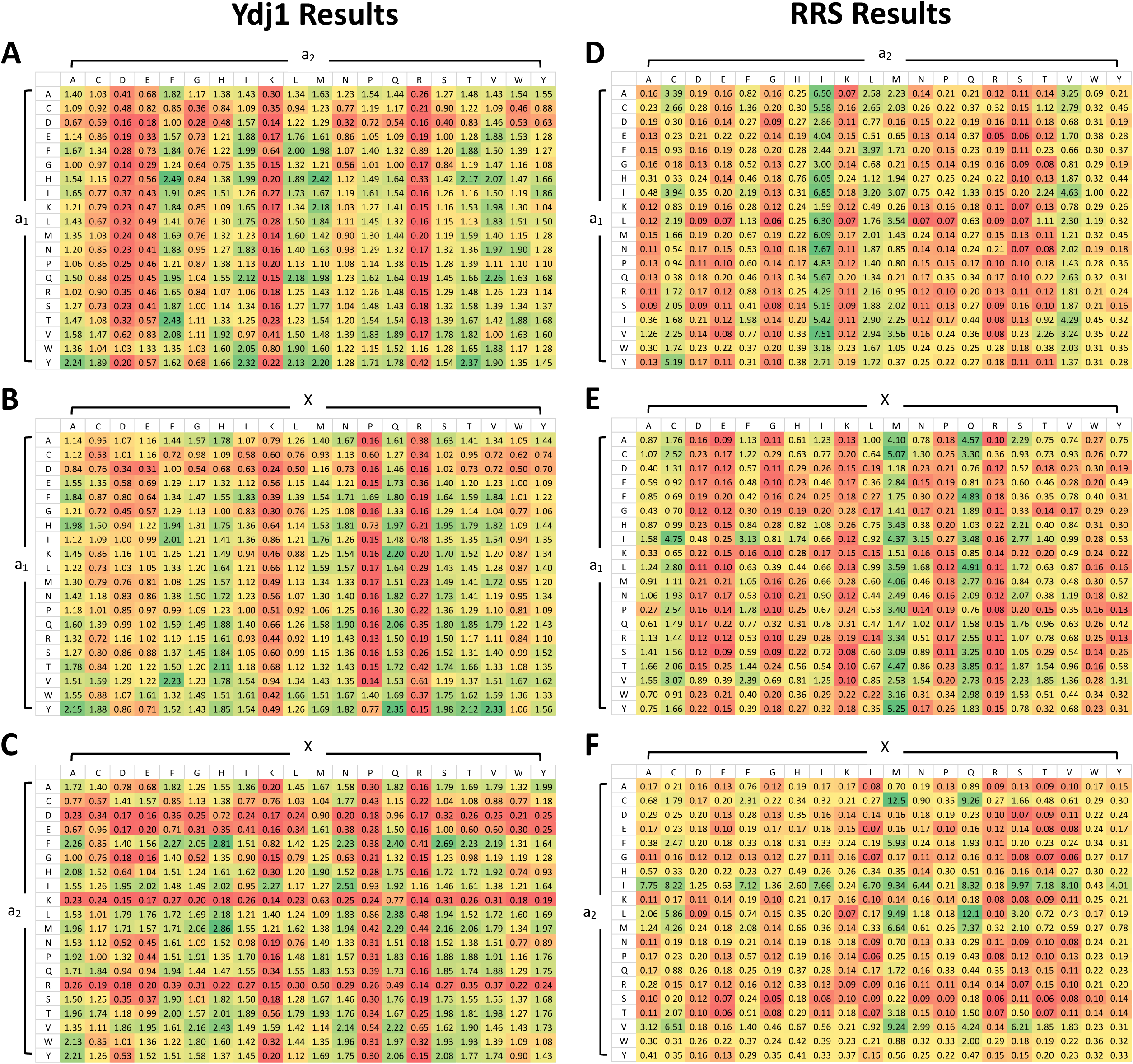
Heatmap analysis reveals contextual determinants of FTase specificity. Ydj1-based comparisons of **A**) a_1_ vs. a_2_, **B**) a_1_ vs. X, and **C**) a_2_ vs. X positions, and Ras-based comparisons of **D**) a_1_ vs. a_2_, **E**) a_1_ vs. X, and **F**) a_2_ vs. X positions. Values were generated by averaging the 20 EFs of each CXXX combinations. Color scheme: red (low likelihood of farnesylation) to green (high likelihood of farnesylation).

For the Ydj1-based data, no positive patterns were observed for the a_1_ position in combination with a_2_ (**Figure 5A**), however, one restrictive pattern, D at a_1_, was observed in combination with X (**Figure 5B**). Generally, a_1_ appears to tolerate many different types of amino acids and does not seem to strongly influence reactivity with FTase in either a positive or negative manner. When considering the a_2_ position, no positive patterns were observed, but several negative patterns were evident. Specifically, D, K and R were restrictive in combination with either a_1_ (**Figure 5A**) or X (**Figure 5C**), with E being additionally restrictive in combination with a_1_. Generally, a_2_ also appears to tolerate many different types of amino acids except for charged amino acids that negatively influence reactivity with FTase. For the X position, a positive pattern, Q at X, was observed in combination with a_1_ (**Figure 5B**). Several negative patterns were also evident at the X position. Both P and R were restrictive in combination with either a_1_ or a_2_, with K being additionally restrictive in the context of a1. Generally, X too seems to tolerate many different types of amino acids except for positively charged amino acids and structurally constrained proline (**Figure 5C**). Thus, despite repeated reports that FTase targets canonical CaaX sequences, our Ydj1-based data strongly indicates that FTase tolerates most CXXX sequences unless amino acids are present that restrict its specificity. Similar heatmap analysis of the Ras-based data also revealed fewer positive than negative patterns. The two positive patterns were I at a_2_ (**Figure 5D**) and M at X (**Figure 5E**), both in combination with a_1_. Most of the negative patterns observed were primarily constrained to a_2_ and X positions in all contexts (**Figure 5C, F**). The a_1_ position, on the other hand, displayed fewest negative patterns; only two patterns were observed at E in combination with a_2_ and K in combination with X. The combined observations that both Ydj1 and Ras-based data reveal high tolerance at the a_1_ position, while a_2_ and X positions have more restrictions is consistent with previous reports on FTase specificity (Reid, Terry et al. 2004, Stein, Kubala et al. 2015).

To fully account for the contextual information from heatmap analysis, the three heatmap values from the Ydj1 NGS dataset were summed for each specific CaaX motif to obtain a heatmap (HM) score. This score represents the sum of 3 averages, where each average accounts for 20 EF data points. The advantage of the HM score is that it normalizes statistical outliers that could skew results when using individual EF scores. The HM scores, when a cutoff of 3 was applied, correlated well with prior evidence of prenylation gathered from previous studies, outperforming published models such as Prenylation Prediction Suite (PrePS), RRS, and our recently published SVM algorithm for predicting farnesylation of non-canonical sequences (Maurer-Stroh and Eisenhaber 2005, Stein, Kubala et al. 2015, Berger, Yeung et al. 2022) (**Table 3**).

**Table 3.** A comprehensive overview of in-house prediction versus published models.

| Protein<sup>1</sup> | Motif (n=48) | Observed<sup>2</sup> | %<sup>2</sup> | PrePS<sup>3</sup> | SVM<sup>4</sup> | RRS EF<sup>5</sup> | YDJ1 HM |
| --- | --- | --- | --- | --- | --- | --- | --- |
| Ras2 | CIIS | + | 100 | 1.281 | 1 | 9.011500 | 4.07 |
| Hmg1 | CIKS | - | 41 | -3.588 | 0 | 0.034717 | 1.93 |
| Rho2 | CIIL | + | 100 | 0.817 | 1 | 10.313800 | 3.65 |
| Ssp2 | CIDL | - | 0 | -4.893 | 0 | 0.254070 | 1.82 |
| Skt5, MiY1 | CVIM | + | 100 | 2.200 | 1 | 3.371400 | 3.66 |
| Tbs1 | CVKM | + | 100 | -2.909 | 0 | 0.066541 | 2.47 |
| YDL022C-A | CSII | + | 100 | 0.987 | 1 | 11.350000 | 3.34 |
| YBR096W | CSEI | - | 0 | -5.759 | 0 | 0.149720 | 1.88 |
| YMR265C | CSNA | + | 100 | -4.848 | 0 | 0.039925 | 3.84 |
| Pet18 | CYNA | + | 100 | -5.843 | 0 | 0.079849 | 4.95 |
| Lih1 | CSGL | - | 61 | -4.577 | 0 | 0.041301 | 2.79 |
| Cup1 | CSGK | - | 59 | -6.461 | 0 | 0.099811 | 1.75 |
| Nap1 | CKQS | + | 100 | -0.975 | 1 | 0.066541 | 4.97 |
| Cst26 | CFIF | + | 100 | -1.924 | 1 | 3.194000 | 4.81 |
| YIL134C-A | CAPY | + | 100 | -2.164 | 1 | 0.199620 | 4.73 |
| Atr1 | CTVA | + | 100 | 0.273 | 1 | 7.775800 | 4.55 |
| Las21 | CALD | + | 100 | -2.887 | 1 | 0.028518 | 4.20 |
| YDL009C | CAVS | + | 100 | 0.237 | 1 | 6.607500 | 4.69 |
| Sua5 | CIQF | + | 100 | -0.601 | 1 | 0.199620 | 5.49 |
| NA | CAAQ | + | 100 | -3.280 | 1 | 0.399250 | 4.82 |
| NA | CAHQ | + | 100 | -3.599 | 1 | 0.399250 | 4.74 |
| NA | CASA | + | 100 | -2.693 | 1 | 0.243980 | 3.91 |
| NA | CKQH | + | 70 | -2.442 | 1 | 0.399250 | 4.37 |
| NA | CNLI | + | 95 | -3.804 | 1 | 0.133080 | 3.84 |
| NA | CSFL | + | 68 | -4.425 | 1 | 0.108890 | 4.28 |
| NA | CVAA | + | 100 | -2.789 | 1 | 0.079849 | 4.81 |
| NA | CVFM | + | 100 | -2.939 | 1 | 2.395500 | 4.76 |
| NA | CKQG | + | 73 | -1.876 | 0 | 0.099811 | 4.06 |
| NA | CKQL | + | 50 | -1.680 | 0 | 0.057035 | 3.84 |
| NA | CQTS | + | 100 | -0.922 | 0 | 0.057035 | 5.44 |
| NA | CQSQ | + | 100 | -1.624 | 1 | 0.159700 | 5.27 |
| NA | CKQA | + | 95 | -1.149 | 1 | 0.133080 | 4.57 |
| NA | CKQC | + | 100 | -0.745 | 1 | 0.099811 | 4.12 |
| NA | CKQD | + | 67 | -4.103 | 0 | 0.066541 | 3.51 |
| NA | CKQE | - | 19 | -3.745 | 0 | 0.066541 | 3.36 |
| NA | CKQF | - | 28 | -2.345 | 0 | 0.057035 | 4.62 |
| NA | CKQH | + | 73 | -2.442 | 1 | 0.399250 | 4.37 |
| NA | CKQI | + | 70 | -1.700 | 1 | 0.099811 | 3.90 |
| NA | CKQK | - | 0 | -4.010 | 0 | 0.036295 | 2.22 |
| NA | CKQM | + | 100 | -1.045 | 1 | 0.133080 | 4.50 |
| NA | CKQN | + | 88 | -5.046 | 0 | 0.199620 | 4.48 |
| NA | CKQP | - | 13 | -2.887 | 0 | 0.399250 | 1.97 |
| NA | CKQQ | + | 100 | -0.747 | 1 | 0.399250 | 5.35 |
| NA | CKQR | - | 0 | -4.533 | 0 | 0.399250 | 1.77 |
| NA | CKQT | + | 96 | -1.874 | 1 | 0.079849 | 4.68 |
| NA | CKQV | + | 100 | -1.744 | 1 | 0.133080 | 4.50 |
| NA | CKQW | + | 71 | -5.184 | 0 | 0.199620 | 3.57 |
| NA | CKQY | + | 100 | -3.326 | 0 | 0.399250 | 4.50 |
| # predicted correctly |  |  |  | 30 | 38 | 17 | 45 |
| # false positive |  |  |  | 0 | 0 | 0 | 2 |
| # false negative |  |  |  | 18 | 10 | 31 | 1 |
| % predicted correctly |  |  |  | 62.5% | 79.2% | 35.4% | 93.8% |
| % false positive |  |  |  | 0.0% | 0.0% | 0.0% | 5.1% |
| % false negative |  |  |  | 64.3% | 50.0% | 75.6% | 11.1% |

<sup>1</sup> Protein or gene name as reported in the Saccharomyces Genome Database
<sup>2</sup> Prenylation status and % gel-shift observed in context of Ydj1
<sup>3</sup> Values reported from (Maurer-Stroh and Eisenhaber 2005); positive cutoff > -2.
<sup>4</sup> Values reported from (Berger, Yeung et al. 2022); positive cutoff > 0.
<sup>5</sup> Values reported from (Stein, Kubala et al. 2015); positive cutoff > 3.

Lastly, we trained a predictive model using the results of the Ydj1-based NGS screen to predict whether a given CXXX sequence can be modified based on a subset of questions (decisions) and possible consequences in a decision-tree model (**Figure 6, Figure S5**). Each question pertains to the whether certain amino acids are present in the a_1_, a_2_, and X positions. The decision tree was created by systematically identifying the best set of questions for predicting whether a CXXX sequence can be modified. This allowed us to identify a general set of rules which govern FTase substrate specificity. Based on our model, the biggest determinant of FTase activity is the restriction of D, E, K, or R at the a_2_ position, followed by the restriction of K, P, or R at the X position. However, amino acids such as I, L, M and V at the a_2_ position seem to neutralize the negative effects of K, P, or R at the X position. Our findings repeatedly raise the likelihood that restrictions, rather than tolerance, are applied to a few impermissible amino acids at a_2_ and X positions of potential substrates modified by FTase. It should be noted that although this simple flowchart is useful in understanding general rules of yeast FTase specificity, HM scores provide a more individualized prediction of the prenylation status of each CXXX sequence.

**Figure 6.**
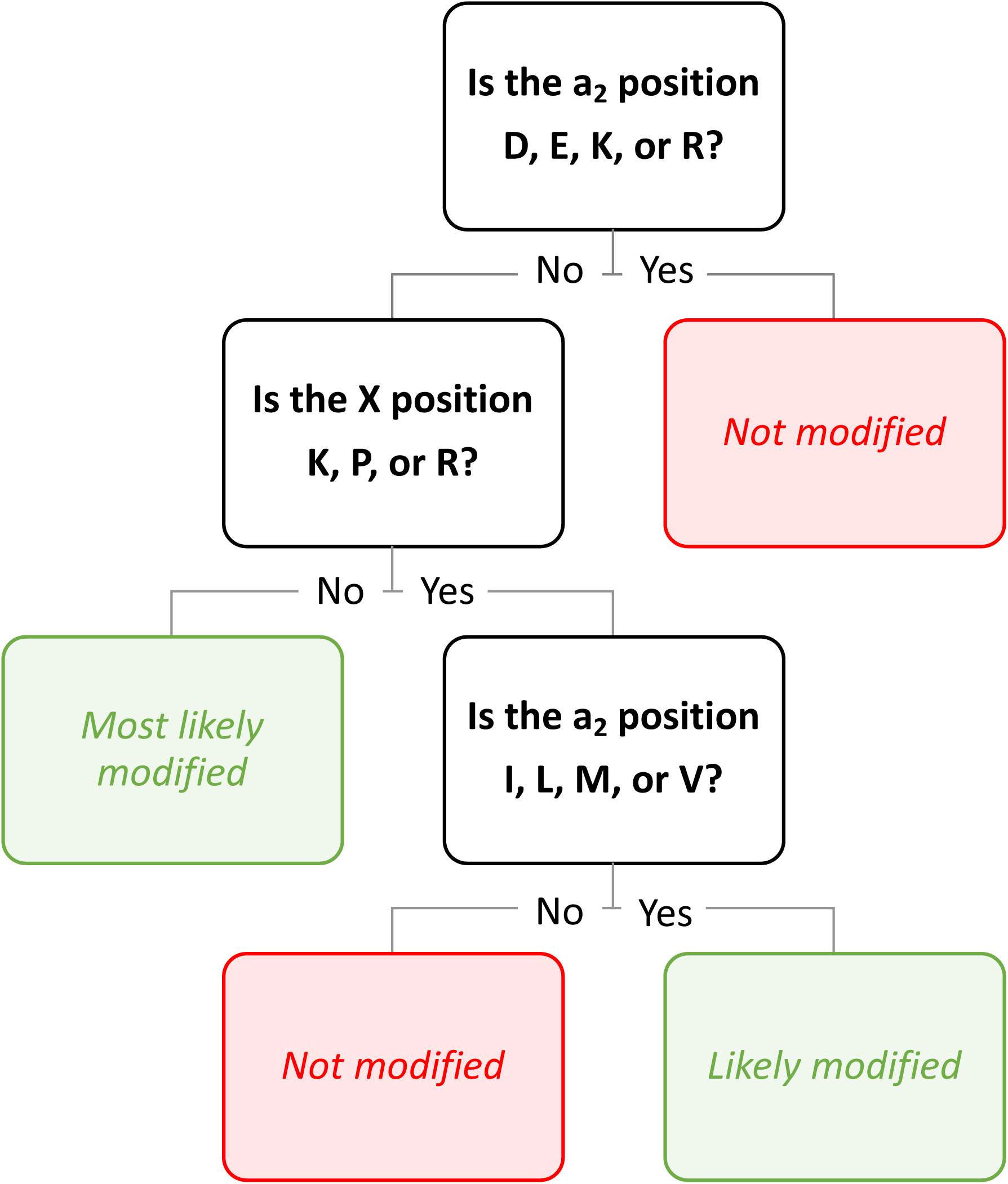
A simplified decision tree for predicting yeast FTase substrates. The output of Ydj1 EF scores was fit into a decision tree model considering each of the variable CaaX motif sites: a_1_, a_2_, and X. Our results strongly suggest that a major determinant of FTase activity depends on a select few discriminatory amino acids at a_2_ (D, E, K, R) and X (K, P, R) positions, with the potential for certain allowable amino acids at a_2_ (I, L, M, V) that can mitigate discrimination caused at the X position. For the full decision tree, see **Figure S5**.

## Discussion

In this study, the use of Ydj1 as a reporter instead of Ras allowed for the identification of yeast FTase targets that extend beyond the canonical CaaX consensus sequence. Notably, this study corroborates previously published observations that non-canonical sequences can be targets of FTase. Restrictions by a small number of amino acids, rather than adherence to the canonical CaaX consensus, seem to guide FTase specificity. We believe that these findings can be used to improve tools for predicting farnesylation status that will outperform current prediction methods.

Structural analysis of mammalian FTase has previously shown that the aaX portion of the protein substrate makes extensive Van der Waals contact with the adjacent isoprenoid group, which is hypothesized to modulate product release (Long, Casey et al. 2002, Reid, Terry et al. 2004). However, these interactions are not predicted when non-aliphatic residues are at a_1_ and a_2_. Our finding that FTase can indeed tolerate non-aliphatic residues at a_1_ and a_2_ raises the possibility that the catalytic site of FTase may display a greater degree of structural flexibility beyond the conformations that could be sampled from mammalian FTase in complex with canonical substrates. Our findings also lead us to predict that binding of charged amino acids (D, E, K, R) may be unstable in the a_2_ binding pocket. Likewise, certain amino acids (K, P, R) at the X position may be unable to coordinate with the product binding pocket of the FTase. These situations may result in the formation of a farnesylated peptide that is unable to be released from the enzyme, thus inhibiting FTase from turning over such substrates efficiently. These possibilities could be resolved by future structural studies involving non-canonical CXXX peptide substrates in complex with either yeast or mammalian FTase, which are structurally and functionally homologous (Kohl, Diehl et al. 1991, Gomez, Goodman et al. 1993, Omer, Kral et al. 1993).

Our study was focused on the recognition of CXXX sequences with respect to FTase specificity. It has been observed, however, that upstream sequences can impact FTase specificity (Cox, Graham et al. 1993). The impact of upstream sequences has been explored by prediction algorithms, most notably by PrePS (Maurer-Stroh and Eisenhaber 2005). It is critical to acknowledge that the Ydj1-CXXX NGS screen recovered sequences that support high temperature growth in the context of Ydj1. Thus, any effect of upstream sequence has not been explored in this work. Additionally, it is possible that some Ydj1-CaaX variants affect growth in ways unrelated to prenylation status, which could impact some of our results. Still, our YDJ1 HM scoring system outperformed predictions made by PrePS, RRS, and SVM machine learning algorithm (Maurer-Stroh and Eisenhaber 2005, Stein, Kubala et al. 2015, Berger, Yeung et al. 2022) (**Table 3**). Combining PrePS predictions that consider upstream sequence in addition to the observations presented in this work is expected to provide the most robust prediction for farnesylation of individual protein targets. As more of the prenylation predictions are validated using additional techniques in the future, it will be possible to determine which prediction methods are most reliable.

Many proteins undergo a process known as isoprenylation. Historically, FTase is recognized as targeting the CaaX sequence but emerging evidence, including that reported here, has indicated that FTase substrates are not limited to the canonical CaaX sequence. Our methods, which primarily focused on yeast FTase, provides insights into the broader specificity of this process which likely extends to human and other FTase enzymes. The discovery of farnesylated non-canonical sequences such as CKQX associated with large families of proteins like Stk11/Lkb1 and Nap1, which lack a canonical CaaX motif, opens a new understanding of post-translational isoprenylation and the potential for future discoveries in this field. Our results also demonstrate that the target specificity of FTase is mainly due to restrictions at the a_2_ and X positions, rather than selection towards a canonical CaaX sequence. As Ras and other canonical reporters rely on multiple modifications for function, previous studies using these proteins may have reported on sequences that are suited for the combined specificities of FTase and downstream enzymes (i.e., CaaX proteases *Sc*Ste24 or *Hs*Rce1) instead of the specificity of FTase alone. Our study, in combination with data from previous studies, provides new ways to advance the understanding of the specificities of each enzymatic step associated with the post-translational isoprenylation pathway, laying out a crucial step toward identifying its cellular targets and the extent to which they are modified.

## Data availability

Yeast strains and plasmids are available upon request. All relevant datasets for this study are included in the supplemental files of the manuscript. **File S1** contains frequency of CXXX sequences observed in naive libraries. **File S2** contains Enrichment Factor results of Ydj1-based NGS. **File S3** contains CKQX hits identified on UniProt. **File S4** contains heatmap analysis. The data derived through the Ras-Recruitment System was previously published and reanalyzed here (Stein, Kubala et al. 2015).

## Abbreviations

FTase: Farnesyltransferase
GGTase: geranylgeranyltransferase
NGS: next-generation sequencing
RRS: Ras Recruitment System
PTM: post-translational modification

## Acknowledgement

We thank Dr. James Hougland (Syracuse University) for early discussions about this work, Dr. Avrom Caplan (City College of New York) for anti-Ydj1 antibody, and Somdut Roy (Georgia Institute of Technology) for the raw Python 3 scripts for the 4D graphical analysis. Lastly, we thank Schmidt lab members for constructive feedback and technical assistance.

## Funding and additional information

This research was supported by Public Health Service grant GM132606 from the National Institute of General Medical Sciences (WKS, NK).

---

**Figure S1.**
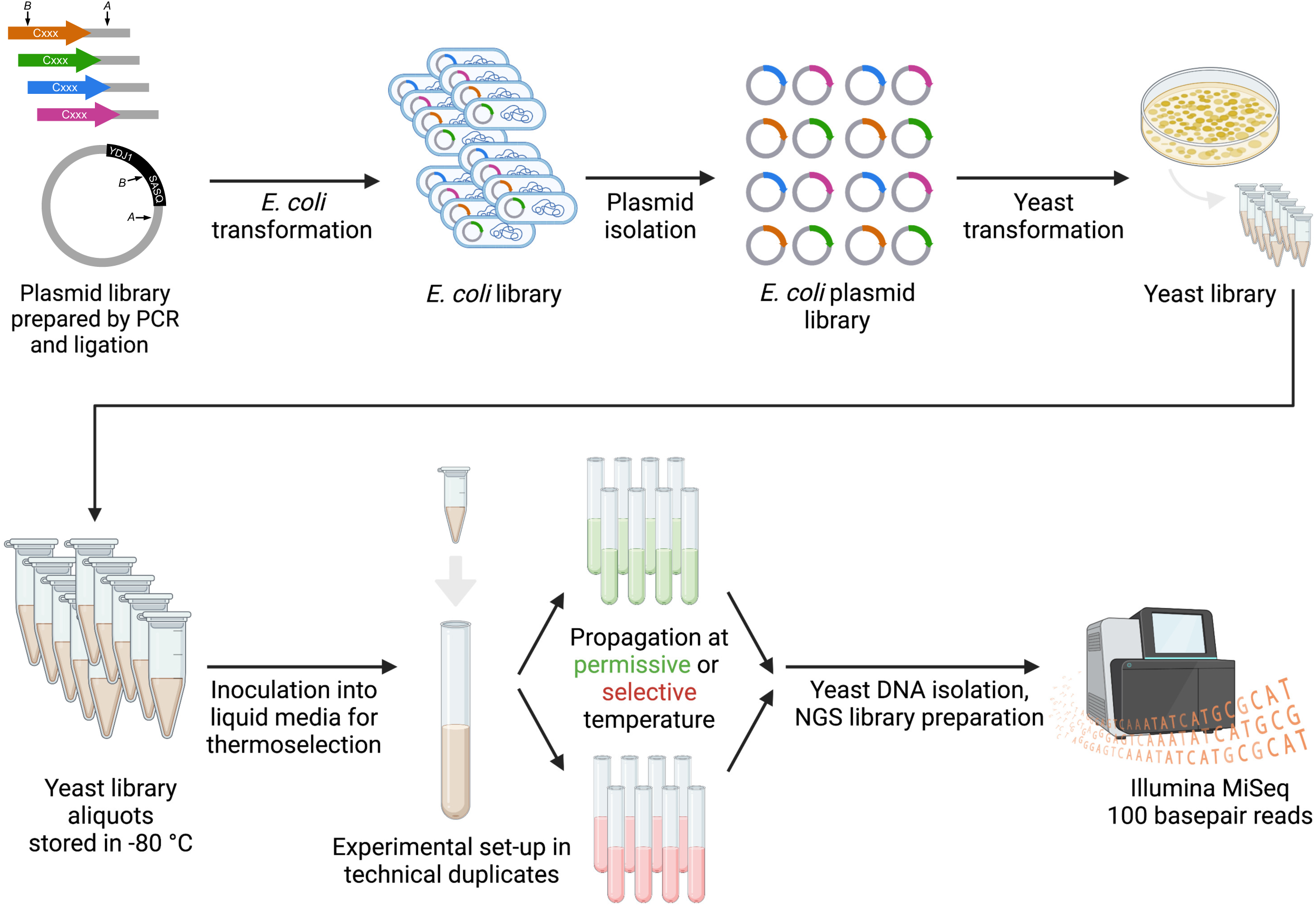
Experimental scheme for genetic screening of Ydj1-CXXX variants assessed via next-generation sequencing (NGS). A plasmid library encoding all 8000 Ydj1-CXXX variants was generated via ligation-based cloning method using PCR fragments derived using Trimer20 oligonucleotides (Integrated DNA Technologies, Newark, NJ). The library DNA was purified from *E. coli*, and plasmids were transformed into the *ydj1Δ* yeast strain. This strain cannot grow at temperatures >37 °C unless complemented by farnesylated Ydj1-CXXX variant. Populations of *ydj1Δ/YDJ1-CXXX* transformants were incubated at permissive (25 °C) and selective (37 °C) temperatures, and plasmids recovered were subject to NGS analysis. Schematic illustration created with Microsoft PowerPoint and BioRender.com.

**Figure S2.**
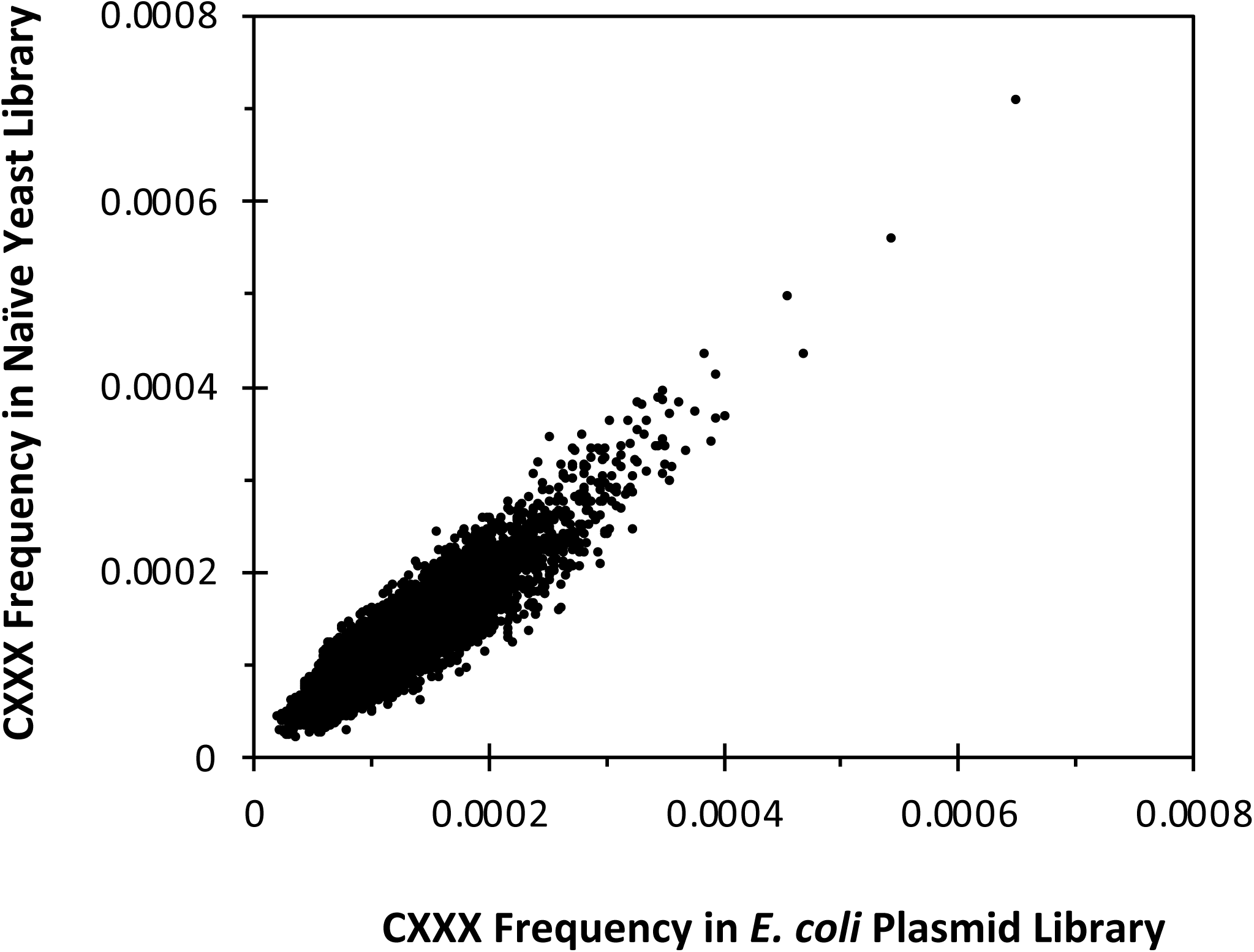
Frequencies of CXXX sequences in naïve yeast library versus E. coli plasmid library. CXXX frequencies from the naïve yeast and *E. coli* plasmid libraries were plotted against each other. No significant change in the frequency profile of individual sequences was observed between the two libraries. Libraries were not perfectly balanced, however, across all CXXX sequences; the extremes exhibited a ∼10x range in relative abundance.

**Figure S3.**
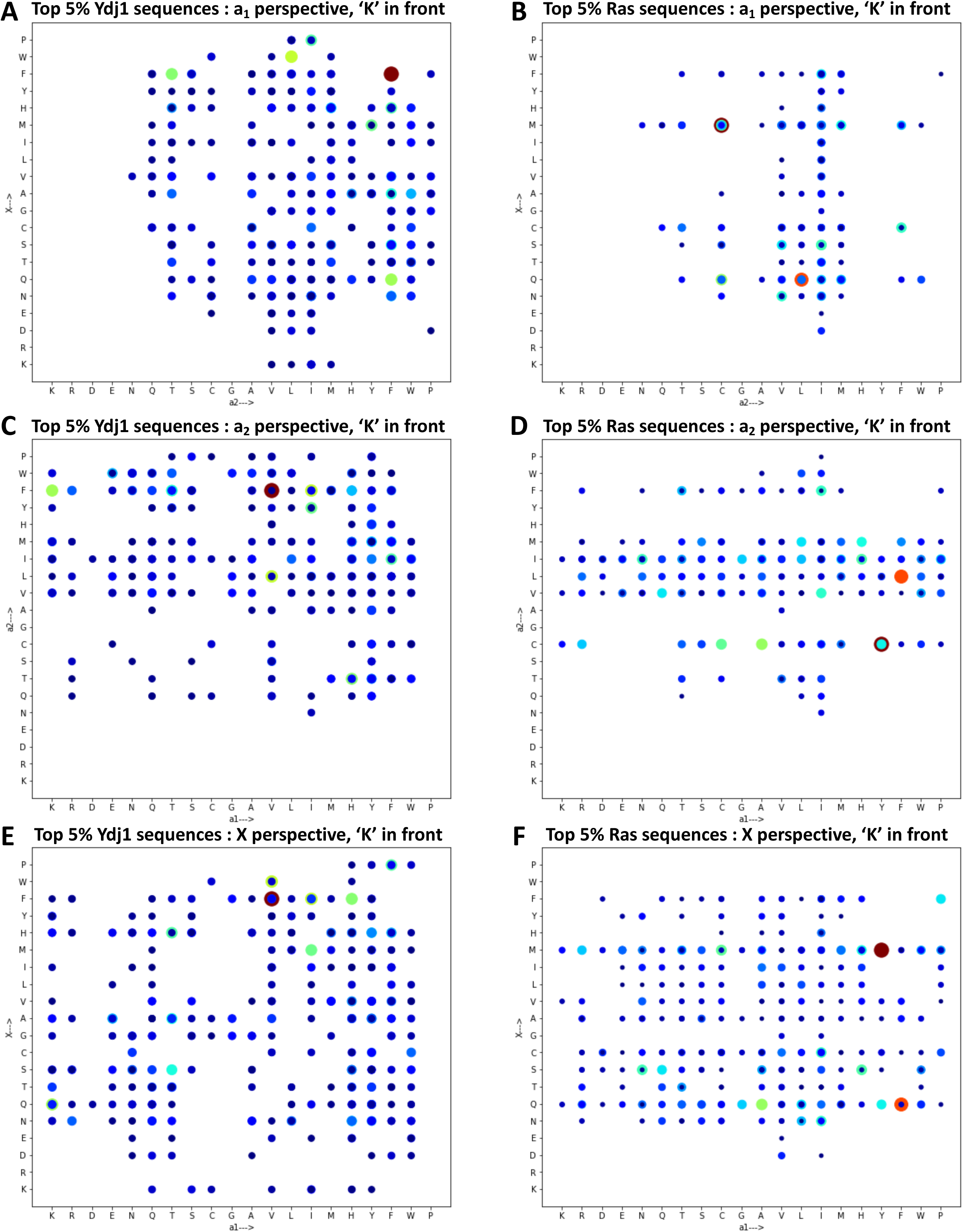
3D analysis of top 5% of sequences from Ydj1-based and Ras-based data sets. The data described in Figures 2E**, F** was plotted in 3D from the perspective of each axis of the 4D plot such that Lysine (K) is the nearest amino acid. The size of the spot is the 3^rd^ dimension and represents relative abundance. The perspectives are along the a_1_ axis (**A**, **B**), the a_2_ axis (**C**, **D**), and X axis (**E**, **F**) for the Ydj1-based data (**A**, **C**, **E**) and the Ras-based data (**B**, **D**, **F**).

**Figure S4.**
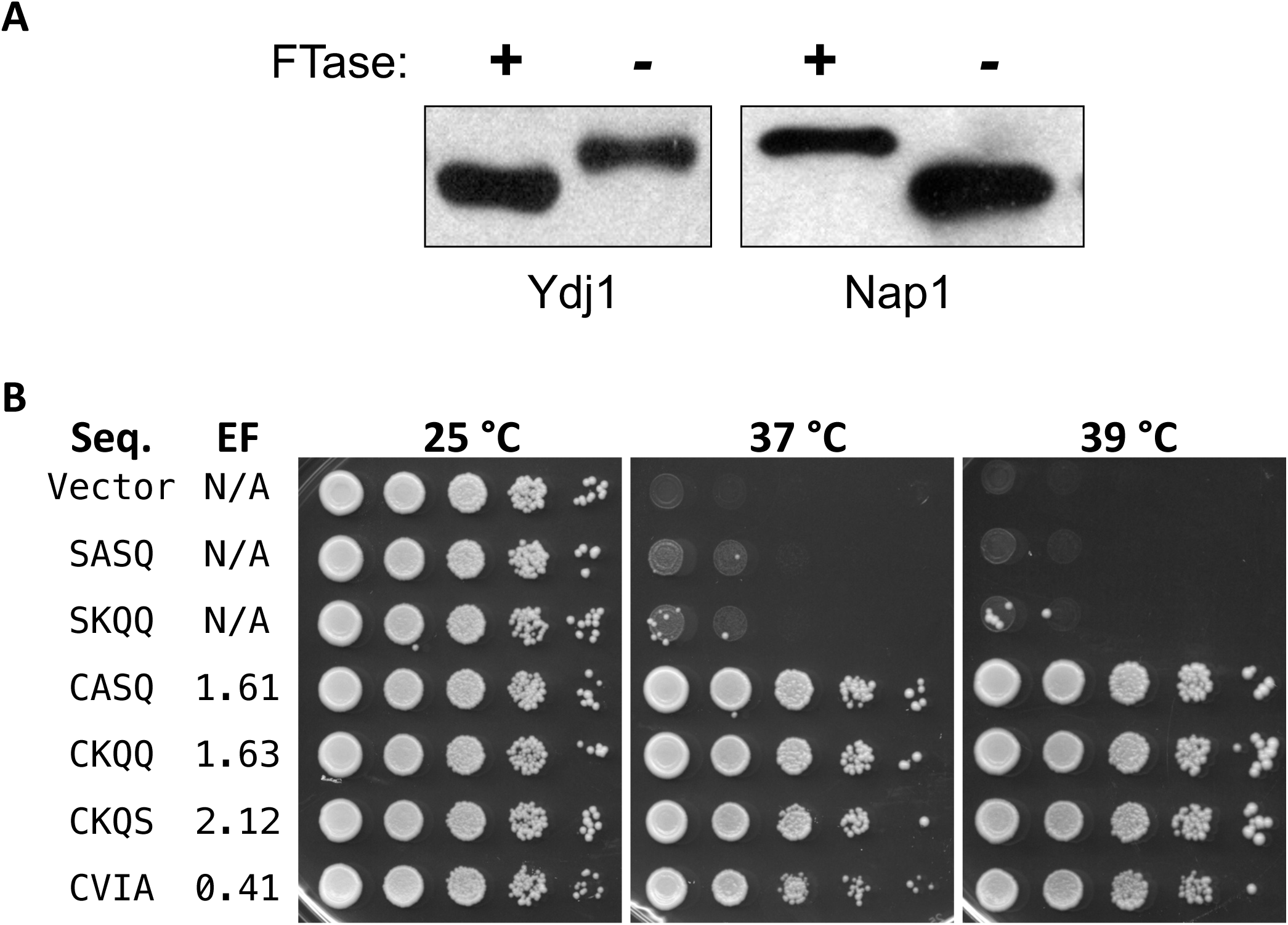
Thermotolerance assay of control sequences. **A**) Gel-shift assays were performed for Ydj1 (CASQ) and His8-Nap1 (CKQS) in the presence and absence of FTase activity. Total yeast extracts were analyzed by SDS-PAGE and immunoblotting. The strains used were BY4741 (+FTase) and yWS1632 (*ram1*Δ; -FTase). **B**) Thermotolerance assays were performed with the indicated Ydj1-CXXX variants as described for Figure 4. Abbreviations: Seq = Sequence, EF = Enrichment Factor, % = percent prenylation, N/A = not applicable.

**Figure S5.**
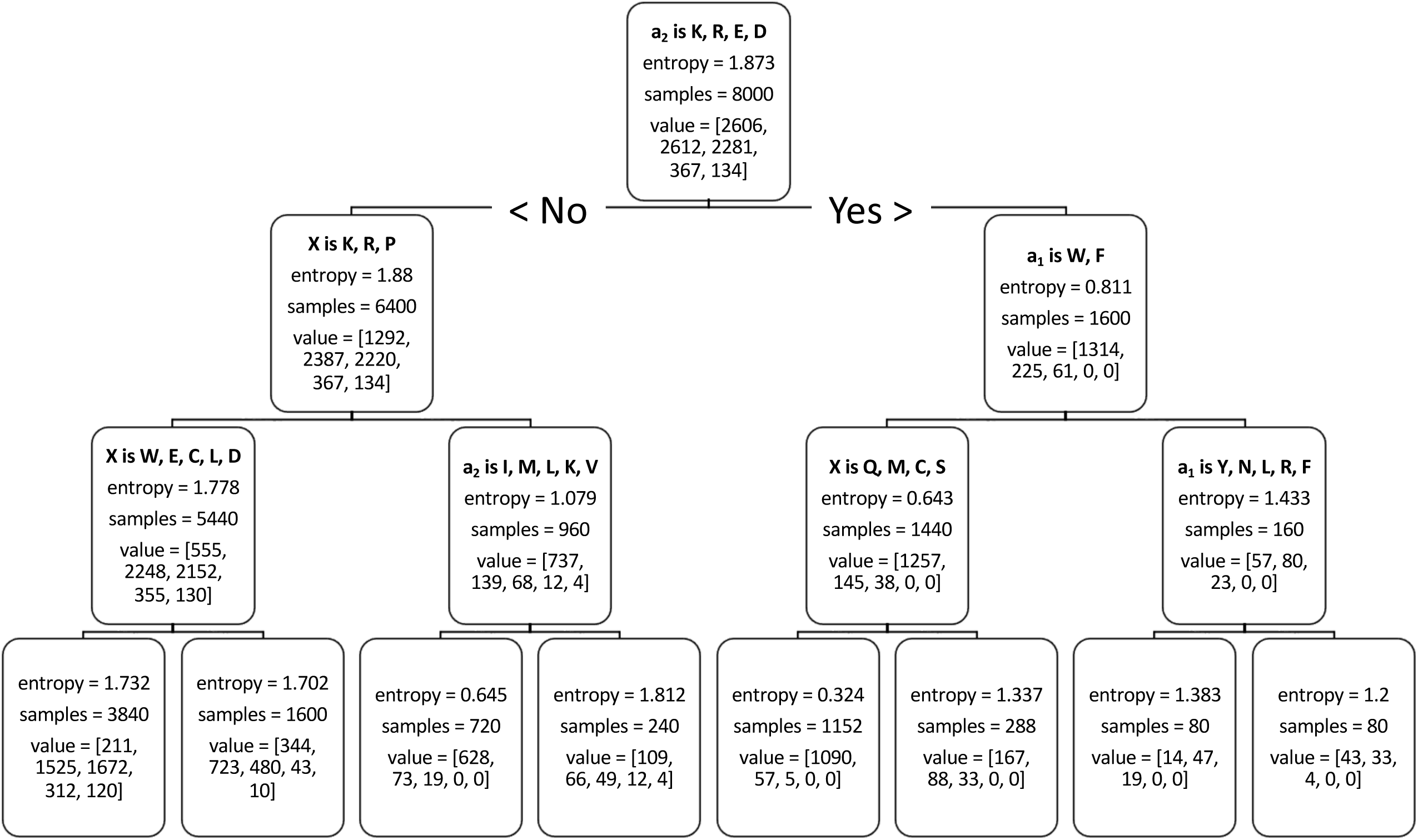
Full decision tree model. Sequence motif features were generated by one-hot encoding each of the variable CaaX motif sites. Additional binary features were included that describe whether the variable residue was within a given set of residues. To define these rules, all possible residue combinations up to 5 amino acids were considered, yielding a total of 65,097 features. While building the decision tree classifier, entropy was used to evaluate the quality of potential splits. Trees were allowed a maximum depth of 3 in order to determine more generalizable rules and all nodes were required to have a minimum of 50 samples. Samples refer to the number of CXXX variants from the previous node that fall within the category. Value refers to the distribution of CXXX variants that fall within specified EF ranges (0-1, 1-2, 2-3, 3-4, >4).

**File S1.** *Frequency of CXXX sequences observed in naive libraries*

**File S2.** *Enrichment factor results of Ydj1-based NGS*

**File S3.** *CKQX hits identified on UniProt*

**File S4.** *Heatmap analysis*

